# Seeding competent TDP-43 persists in human patient and mouse muscle

**DOI:** 10.1101/2024.04.03.587918

**Authors:** Eileen M. Lynch, Sara Pittman, Jil Daw, Chiseko Ikenaga, Sheng Chen, Dhruva D. Dhavale, Meredith E. Jackrel, Yuna M. Ayala, Paul Kotzbauer, Cindy V. Ly, Alan Pestronk, Thomas E. Lloyd, Conrad C. Weihl

## Abstract

TAR DNA-binding protein 43 (TDP-43) is an RNA binding protein that accumulates as aggregates in the central nervous system of some neurodegenerative diseases. However, TDP-43 aggregation is also a sensitive and specific pathologic feature found in a family of degenerative muscle diseases termed inclusion body myopathy (IBM). TDP-43 aggregates from ALS and FTD brain lysates may serve as self-templating aggregate seeds *in vitro* and *in vivo,* supporting a prion-like spread from cell to cell. Whether a similar process occurs in IBM patient muscle is not clear. We developed a mouse model of inducible, muscle-specific cytoplasmic localized TDP-43. These mice develop muscle weakness with robust accumulation of insoluble and phosphorylated sarcoplasmic TDP-43, leading to eosinophilic inclusions, altered proteostasis and changes in TDP-43-related RNA processing that resolve with the removal of doxycycline. Skeletal muscle lysates from these mice also have seeding competent TDP-43, as determined by a FRET-based biosensor, that persists for weeks upon resolution of TDP-43 aggregate pathology. Human muscle biopsies with TDP-43 pathology also contain TDP-43 aggregate seeds. Using lysates from muscle biopsies of patients with IBM, IMNM and ALS we found that TDP-43 seeding capacity was specific to IBM. Surprisingly, TDP-43 seeding capacity anti-correlated with TDP-43 aggregate and vacuole abundance. These data support that TDP-43 aggregate seeds are present in IBM skeletal muscle and represent a unique TDP-43 pathogenic species not previously appreciated in human muscle disease.

**Summary:** TDP-43 aggregate seeds persist in mouse and human skeletal muscle independent of large TDP-43 inclusions.

## INTRODUCTION

TAR DNA-binding protein-43 (TDP-43) is a ubiquitously expressed RNA-binding protein which is primarily localized to cell nuclei. However, certain disease conditions cause TDP-43 to form cytoplasmic aggregates, resulting in a loss of function through interruption of RNA processing and potentially causing a toxic gain of function as well. TDP-43 aggregation has been most commonly studied in the context of neurodegenerative diseases such as amyotrophic lateral sclerosis (ALS) and frontotemporal dementia (FTD). Within neurodegeneration, TDP-43 aggregates behave similarly to aggregates of tau and α-synuclein which have prion-like properties. These misfolded prion-like proteins form insoluble aggregates and are able to induce their respective monomers to misfold and aggregate, a process referred to as seeding. These proteins have been shown to propagate to other brain regions, suggesting trans-synaptic spread through neuronal networks [1]. Experimentally, mouse brains injected with the extract from brains of patients with frontotemporal lobar degeneration (FTLD) show a progressive expansion of TDP-43 pathology into neighboring regions, supporting the concept of cell-to-cell transmission through neuronal networks [2, 3]. This correlates with human brain studies of ALS, FTD, Alzheimer’s disease and limbic-predominant age-related TDP-43 encephalopathy (LATE) patients which show a specific pattern of TDP-43 pathology progression through brain regions [4–7].

In addition to its role in neuronal pathology, TDP-43 has also been found in sarcoplasmic aggregates from the skeletal muscle of patients with several protein aggregate myopathies, particularly sporadic inclusion body myositis and inherited inclusion body myopathy [8–16]. While mutations in *TARDBP* can cause ALS and FTLD, until recently there were no *TARDBP* mutations linked to pure myopathy. However, several new studies have reported on four families with a rimmed vacuolar myopathy and IBM-like pathology characterized by TDP-43 aggregates. These families were found to have dominantly inherited missense or frameshift mutations in *TARDBP* [17, 18]. The discovery of these causative mutations supports the significance of TDP-43 aggregate pathogenicity in skeletal muscle. Additionally, skeletal muscle is known to incubate prion protein in diseases such as bovine spongiform encephalopathy (BSE), scrapie, and chronic wasting disease [19–21]. However, whether TDP-43 aggregates have the capacity to spread along the length of a myofiber or from myofiber to myofiber and whether skeletal muscle can incubate TDP-43 aggregate seeds akin to prion diseases has not been established.

TDP-43 forms insoluble sarcoplasmic TDP-43 aggregates termed myo-granules during normal skeletal muscle differentiation and regeneration. These myo-granules organize with the mRNA of skeletal muscle structural proteins, facilitating the timely translation of these proteins [22]. We hypothesize that the physiologic process of muscle regeneration and differentiation in skeletal muscle creates insoluble TDP-43 myo-granules that resolve in healthy patients but may persist in patients that develop IBM or other myopathies with TDP-43 aggregate pathology. This may be due to pathogenic variants in TDP-43 or variations in other proteins known to be associated with TDP-43 aggregation and IBM such as valosin containing protein (VCP) and SQSTM1 [23–25].

The detection of these TDP-43 aggregate seeds may serve as a marker of disease pathogenesis and be a tractable therapeutic target aimed at the insidious progression of disease pathology from fiber to fiber.

## RESULTS

### TDP-43 seeding is present in patient tissue with TDP-43 aggregate pathology

To detect TDP-43 aggregate seeds from tissue of patients with TDP-43 proteinopathy, we used a previously described TDP-43 FRET biosensor line [26]. The TDP-43 FRET biosensors were generated via stable expression of two independently tagged human TDP-43 C-terminal fragments (mClover or mRuby) in HEK293 cell lines. The exogenous application of lipofectamine-encapsulated TDP-43 seeds to the biosensors templates the aggregate conversion of the mClover and mRuby tagged C-terminal fragments, allowing FRET to occur **(Figure 1A)**. FRET signal is detected via flow cytometry and quantitated as integrated FRET density (percent FRET positive cells x FRET signal intensity). TDP-43 biosensor specificity for TDP-43 seeds was demonstrated by the application of recombinant monomeric TDP-43 and TDP-43 pre-formed fibrils (PFF). As previously demonstrated, TDP-43 PFF but not monomeric TDP-43 induced FRET in a concentration-dependent manner (**Figure 1B-C**).

**Fig. 1.**
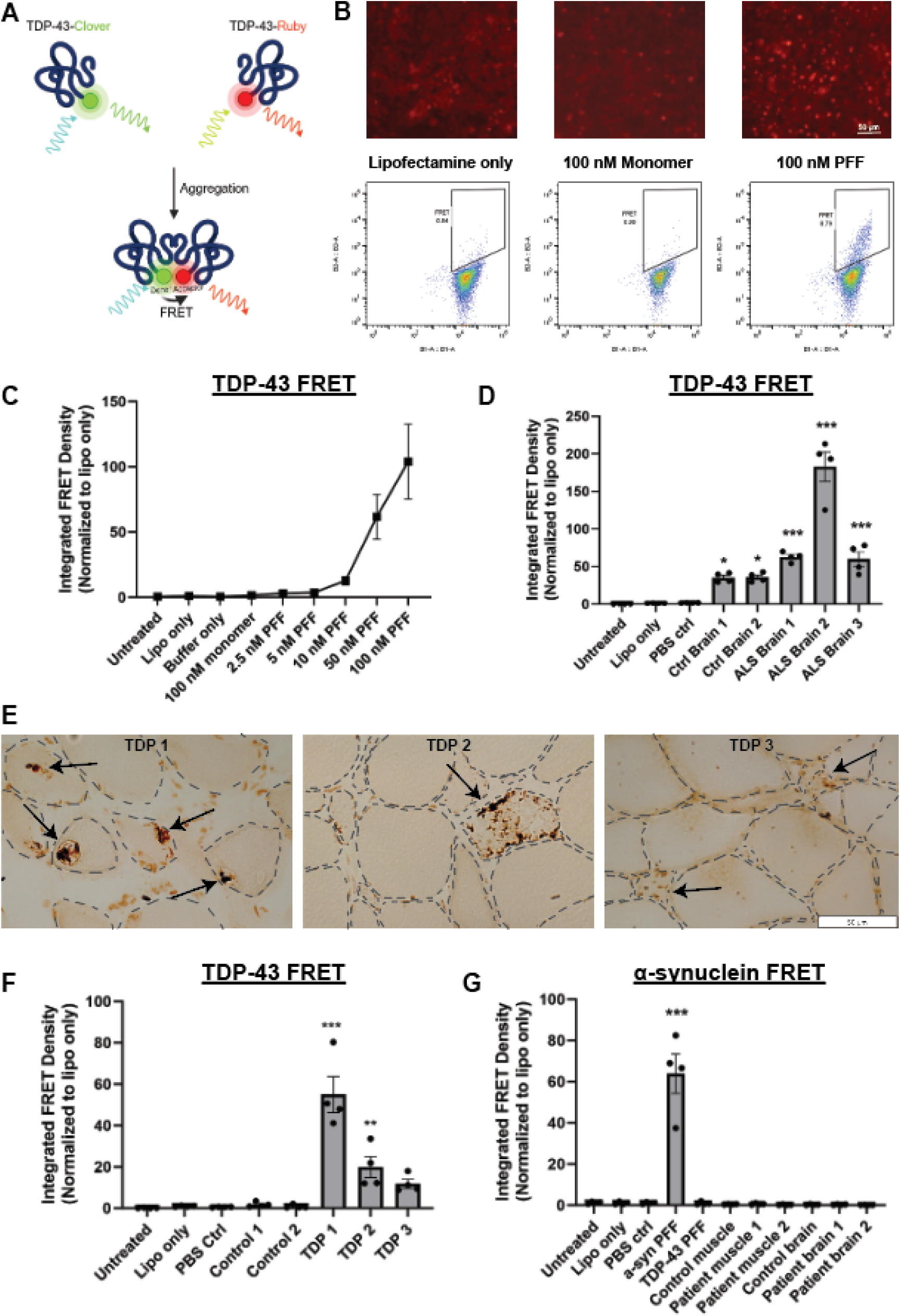
Confirmation of TDP-43 seeding in patient brain and muscle samples using a FRET sensor line. (**A**) TDP-43 FRET biosensors express two independently tagged c-terminal fragments (aa262-414) of human TDP-43 fused to mruby and mclover. When TDP-43 aggregation occurs, the two fluorophores are close enough for an energy transfer to occur and the signal is detected by flow cytometry. (**B**) Representative fluorescent images of the FRET sensor line and the flow cytometry gating strategy using lipofectamine alone, 100nM TDP-43 monomeric protein, or 100 nM TDP-43 preformed fibrils (PFF). Note only TDP-43 PFF induces a FRET signal. (**C**) A dose response curve of the TDP-43 biosensor line treated with buffer, lipofectamine, TDP-43 monomer, or increasing concentrations of recombinant TDP-43 PFF. (**D**) The RIPA-insoluble fraction of human ALS patient autopsy brain tissue was added to the FRET sensor line with lipofectamine and analyzed by flow cytometry after 72 hours. Note the increase in FRET signal from ALS brain tissue. (**E**) Representative immunohistochemical images of TDP-43 staining from human muscle biopsies with TDP-43 inclusions (arrows). Black dotted lines outline individual muscle fibers. (**F**) The RIPA-insoluble fraction from control muscle or muscle biopsies confirmed to have positive TDP-43 aggregate staining were added to the FRET sensor line to confirm seeding. (**G**) Samples from muscle and brain lysates with established TDP-43 seeding ability were added to an α-synuclein FRET sensor line to test for TDP-43 seeding specificity. In contrast to TDP-43 biosensors, no α-synuclein seeding is present. FRET signal is measured as integrated FRET density, % positive cells x median fluorescence intensity and normalized to a lipofectamine only control. *P<0.05, **P<0.01, ***P<0.001 by one-way ANOVA followed by Dunnett’s multiple comparisons test to lipofectamine control.

To identify TDP-43 aggregate seeds from human tissue with TDP-43 proteinopathies, we homogenized human postmortem motor cortex samples from two control and three ALS patients with TDP-43 pathology in RIPA buffer to generate an insoluble fraction. The insoluble fraction was resuspended in PBS and delivered to the TDP-43 biosensors using lipofectamine (**Figure 1D**). ALS brain lysates had an increased integrated FRET density when compared with control brain lysates. To evaluate TDP-43 seeding from muscle-derived TDP-43 proteinopathies, we performed a similar experiment using the insoluble lysates from skeletal muscle biopsies of two normal control muscles and three muscle biopsies with TDP-43 pathology (**Figure 1E**). The insoluble fraction from these biopsy lysates also seeded TDP-43 aggregation in the TDP-43 biosensors (**Figure 1F**). To establish whether the induction of TDP-43 aggregation was specific to TDP-43 or whether this was due to a non-specific induction of aggregation, we similarly added patient muscle and brain lysates that had TDP-43 seeding capacity to an established α-synuclein biosensor cell line [27]. While α-synuclein PFF induced FRET in the α-synuclein biosensors, TDP-43 PFF and muscle or brain from TDP-43 proteinopathy patients did not (**Figure 1G**).

### Overexpression of cytoplasmic-mislocalized TDP-43 recapitulates TDP-43 myopathology in mouse skeletal muscle

To understand the relationship between TDP-43 aggregation, myopathology, and TDP-43 seeding in skeletal muscle, we developed an inducible mouse model of muscle-specific sarcoplasmic TDP-43 aggregation. These mice express human TDP-43 with a mutated nuclear localization signal specifically in skeletal muscle under the human skeletal actin promoter (HSA-hTDP43^ΔNLS^) using a tetracycline-controlled conditional expression system similar to those used for the CamKIIa-hTDP43^ΔNLS^ mice and rNLS8 mice [28, 29]. To do this, we crossed tetO-hTDP43^ΔNLS^ with HSA-rtTA mice. The resulting HSA-hTDP43^ΔNLS^ mice only express the transgene upon activation with doxycycline (dox) chow diet (**Figure 2A**). This allows temporal control of transgene expression. Over the course of 4 weeks of dox treatment, mice experienced progressive weight loss, loss of motor function, and ultimately death at or shortly after 4 weeks of transgene expression (**Figure 2B**). Mice switched back to regular chow at the 4 week timepoint were unable to recover and survived no longer than 1 week following the 4 weeks of treatment with dox chow. Muscle histopathology demonstrated varied fiber size with frequent smaller angular fibers following 4 weeks of doxycycline treatment. Most fibers had subsarcolemmal eosinophilic inclusions often adjacent to myonuclei on hematoxylin and eosin staining. NADH staining highlighted normal internal architecture with the exception of subsarcolemmal clearings consistent with the presence of aggregated protein (**Figure 2C**). Hind limb muscle weights and overall body weights were significantly decreased compared to age-matched untreated control mice (**Figure 2D**). By 3 weeks of dox treatment, kyphosis of the spine was noted, consistent with global muscle weakness. X-rays were taken to calculate the kyphosis index which was significant in mice treated with dox for 3 weeks compared to age-matched controls (**Figure 2E**).

**Fig. 2.**
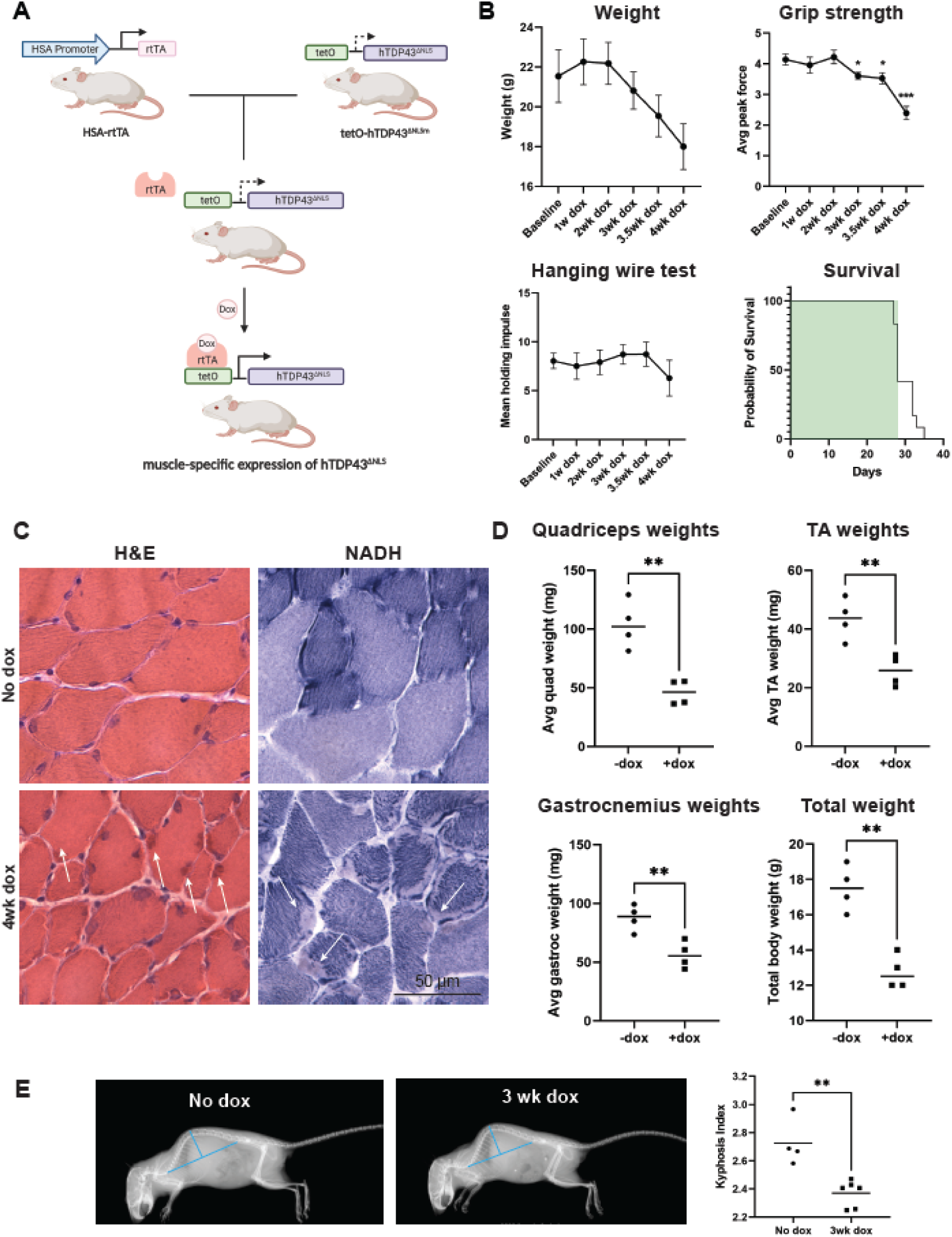
Mice expressing cytoplasmic hTDP-43 develop a myopathy with premature death. (**A**) HSA-rtTA mice were crossed with tetO-hTDP-43^ΔNLS^ mice to create doxycycline-inducible muscle-specific expression of hTDP-43^ΔNLS^. (**B**) A cohort of mice were studied over the course of a 4 week dox treatment (shaded in green on the survival curve), tracking total body weight, strength, and survival time. N = 13 for survival curve, N = 11 for weight and grip strength, N = 7 for hanging wire as 4 mice were too frail to perform the test. (**C**) After 4 weeks of dox treatment, H&E staining showed mild myopathic changes with varied fiber sizes. Non-nuclear eosinophilic staining in H&E and pale regions on NADH staining (arrows) highlight sarcoplasmic inclusion. (**D**) Mice treated with dox chow for 4 weeks compared to age-matched no dox controls showed decreased hind limb muscle weights as well as decreased total body weight compared to untreated controls. (**E**) Example x-ray images of spinal kyphosis in HSA-hTDP-43^ΔNLS^ mice after 3 weeks on dox chow and the calculated kyphosis index. *P>0.05, **P<0.01, ***P<0.001; unpaired Student’s t-test.

### HSA-hTDP43^ΔNLS^ mice develop sarcoplasmic accumulation of insoluble TDP-43 aggregates

To explore the progression of TDP-43 aggregate formation, mouse hindlimb (gastrocnemius) muscles were collected for analysis following 2 weeks and 4 weeks of dox-induced transgene activation. Immunohistochemistry showed abundant subsarcolemmal TDP-43 aggregates that corresponded to phospho-TDP-43 staining which is often perinuclear in localization (**Figure 3A**). Insoluble fractionation followed by western blot analysis showed progressive accumulation of RIPA-insoluble TDP-43/pTDP-43 over time (**Figure 3B**). Given that the mice die around 4 weeks of transgene expression but without major structural changes to the skeletal muscles, TDP-43 transgene expression was examined in other organ systems. Indeed, there is increased total TDP-43 in the heart with activation of the transgene to a similar level of that found in hindlimb muscles, contributing to mortality (**Figure 3C**). Changes in the level of proteins such as p62, HSP70, and lamp2 indicate a loss of proteostasis over time (**Figure 3D**). Additionally, pTDP-43 aggregates co-localized with p62 and ubiquitin (**Figure 3E**).

**Fig. 3.**
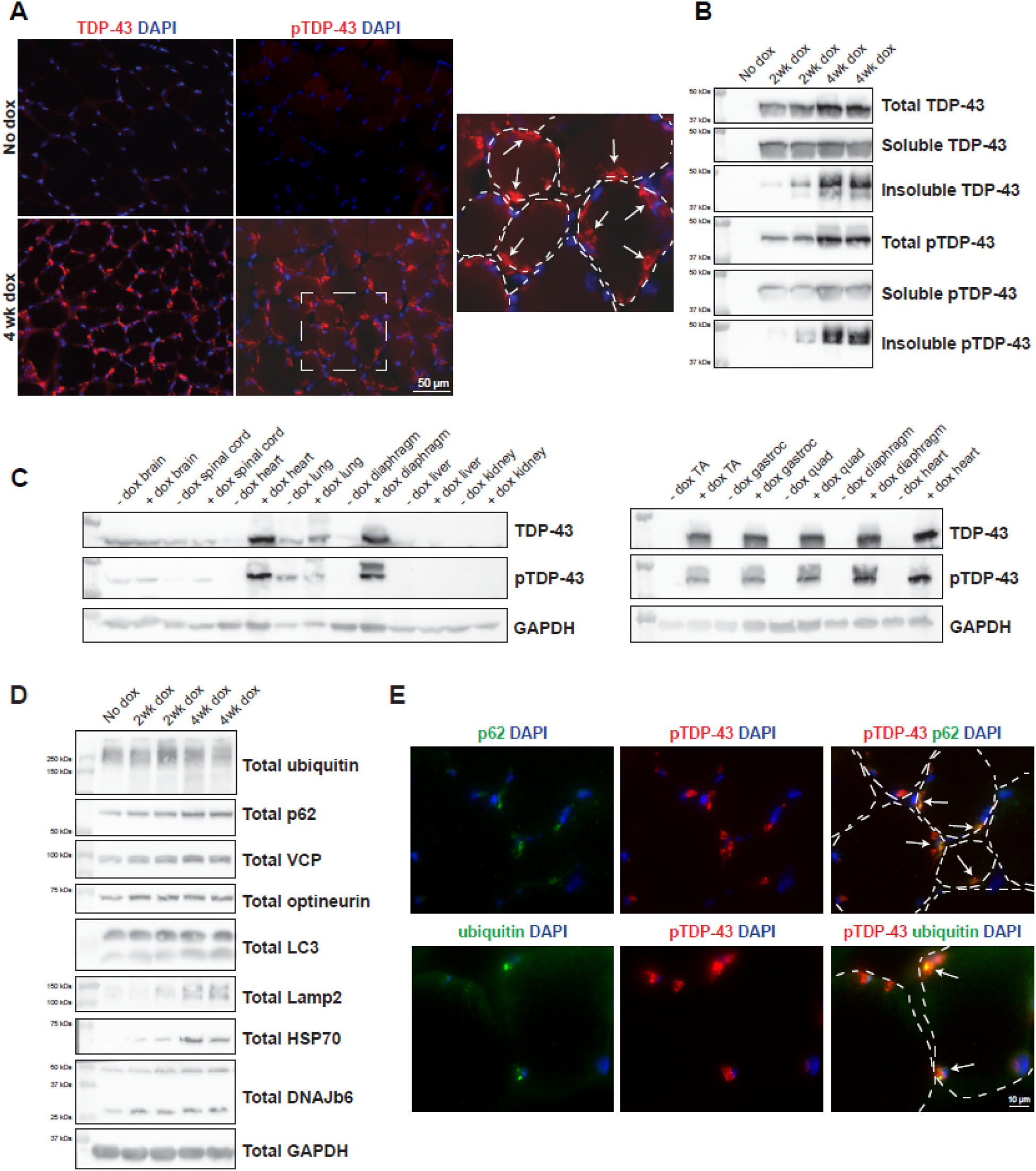
HSA-hTDP-43^ΔNLS^ mice develop abundant sarcoplasmic insoluble phosphorylated TDP-43 aggregates which disrupt proteostasis. (**A**) Representative immunofluorescent images of hindlimb cross sections of HSA-hTDP-43^ΔNLS^ mice after 4 weeks of transgene activation by dox chow, showing abundant TDP-43 (red upper panel) and phosphorylated TDP-43 aggregates (red lower panel). Dashed white lines indicate muscle fiber outlines. (**B**) Gastrocnemius muscle lysates from HSA-hTDP-43^ΔNLS^ mice were processed for insoluble fractionation followed by western blot to detect total, soluble, and insoluble levels of TDP-43 and pTDP-43 which showed an increase over time. The total GAPDH can be found in part **D**. (**C**) Mice treated with dox chow for 4 weeks were evaluated for TDP-43 transgene expression in non-skeletal muscle tissue. Notably, TDP-43 was increased in cardiac tissue. (**D**) Gastrocnemius muscle lysates from HSA-hTDP-43^ΔNLS^ mice processed at the indicated time points were evaluated using antibodies against ubiquitin, p62, VCP, optineurin, LC3, Lamp2, HSP70, DNAJB6, and GAPDH. (**E**) Representative dual fluorescent imaging of pTDP-43 (red) and p62 (green upper panel) or ubiquitin (green lower panel) at 2 weeks of dox treatment. Dashed white lines indicate muscle fiber outlines.

### TDP-43-related pathology is resolved by three weeks following transgene suppression

To understand whether insoluble TDP-43 aggregates persisted or were cleared in skeletal muscle following removal of doxycycline, we fed HSA-hTDP43^ΔNLS^ mice with dox chow to activate the transgene for 2 weeks, then returned to regular chow to allow for recovery. Surprisingly, following 3 weeks of recovery, sarcoplasmic TDP-43 aggregates are absent as demonstrated by immunohistochemistry for TDP-43 and pTDP-43 of hindlimb muscles (**Figure 4A**). Insoluble TDP-43 as detected by fractionation western blot is also absent following 3 weeks of recovery (**Figure 4B**). Markers of proteostatic stress (HSP70 and high molecular weight ubiquitin) increased at two weeks and then similarly resolved following three weeks of recovery (**Figure 4C**). Notably, following three weeks recovery, skeletal muscle pathology has small fibers with centralized nuclei and an increase in connective tissue (**Figure 4D**).

**Fig. 4.**
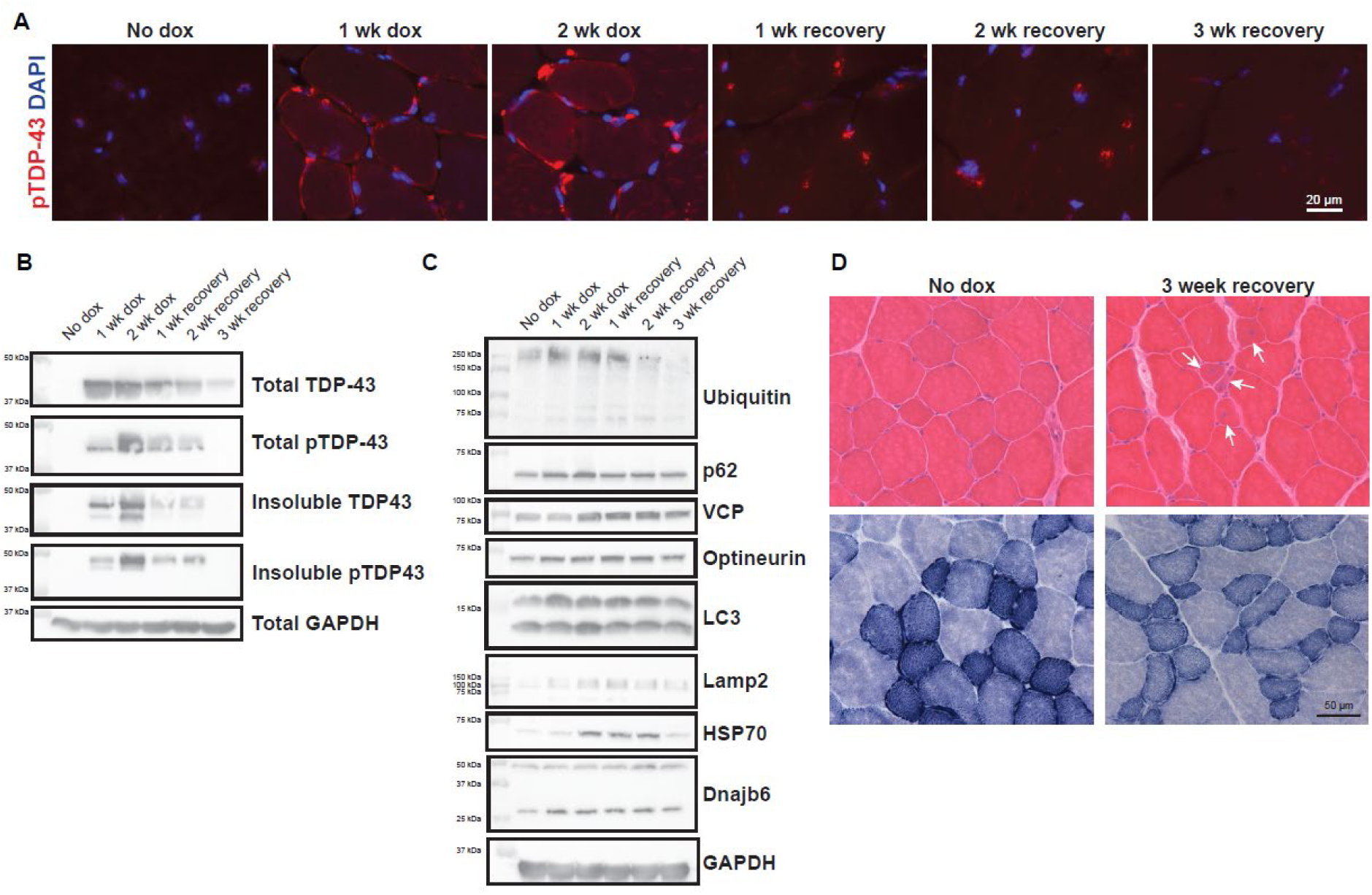
TDP-43 aggregates are cleared and proteostasis resolved following removal of doxycycline. (**A**) Representative immunofluorescent images of quadriceps femoris muscles stained for pTDP-43 (red) in HSA-hTDP43^ΔNLS^ mice at the indicated timepoints when treated with dox chow for 2 weeks followed by a return to a regular chow diet to turn off transgene expression. By three weeks no pTDP-43 is seen. (**B**) Insoluble fractionation western blot of HSA-hTDP43^ΔNLS^ mouse gastrocnemius muscles shows accumulation and then resolution of insoluble TDP-43 and pTDP-43 over the time course of two weeks doxycycline followed by 3 weeks of recovery. (**C**) Gastrocnemius muscle lysates from HSA-hTDP43^ΔNLS^ mice processed at the indicated time points were evaluated using antibodies against ubiquitin, p62, VCP, optineurin, LC3, Lamp2, HSP70, DNAJB6 and GAPDH. (**D**) Representative H&E and NADH images of quadriceps femoris muscle after 3 weeks of recovery as compared to no dox controls. Note the presence of smaller fibers with centralized nuclei (arrows).

Transmission electron microscopy of tibialis anterior muscle following 2 weeks of dox treatment shows granular accumulations corresponding to the localization of TDP-43 aggregates by prior immunostaining. Accordingly, the aggregates are subsarcolemmal and often adjacent to nuclei (**Figure 5B-C**). Aggregates have a granular structure associated with scattered fibrils (**Figure 5D**). Upon removal of doxycycline, aggregates resolve and are often infiltrated with vesicular and vacuolar structures (**Figure 5E-F**).

**Fig. 5.**
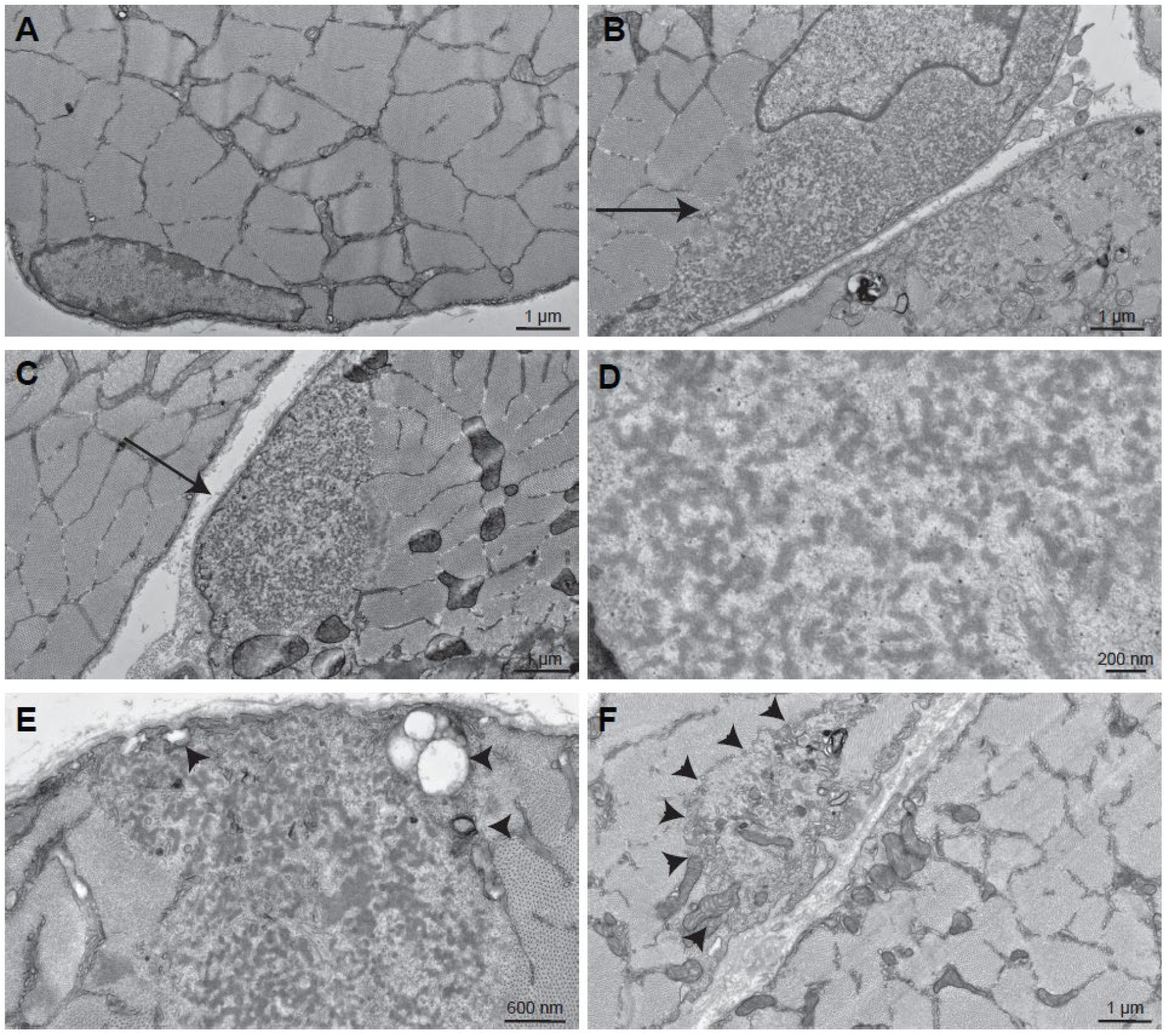
Ultrastructural analysis of HSA-hTDP-43^ΔNLS^ mouse muscle. (**A**) Control TA muscle fiber with myonuclei. (**B-D**) HSA-hTDP-43^ΔNLS^ mice treated for two weeks with doxycycline. Note large granular amorphous inclusions that are subsarcolemmal and myonuclei-adjacent (arrows). (**D**) Higher magnification of the granular structure of an aggregate from a mouse on dox treatment for 2 weeks. (**E-F**) HSA-hTDP-43^ΔNLS^ mice treated for two weeks with doxycycline and then changed to normal chow for three weeks. Note autophagic debris and vacuolation in and around the inclusion (arrowheads).

One marker of TDP-43 dysfunction in tissue is alterations in RNA splicing. In particular, the inclusion of select cryptic exons suggests a loss of TDP-43 functionality. To determine if there were changes in splicing related to dysregulation of normal TDP-43 and RNA interactions, we evaluated hindlimb muscle for mouse muscle-specific cryptic exon marker in the gene *Sh3gbr* as reported by Jeong et al [30]. More slowly migrating bands indicate a 45 base pair cryptic exon inclusion between exons 1 and 2 of *Sh3gbr*. Cryptic exon inclusion was apparent following one or two weeks on dox but disappeared following an additional week of recovery (**Figure 6B-C**). Surprisingly, the cryptic exon inclusion resolved past two weeks on dox even if the dox treatment was continued out to 4 weeks (**Figure 6D**).

**Fig. 6.**
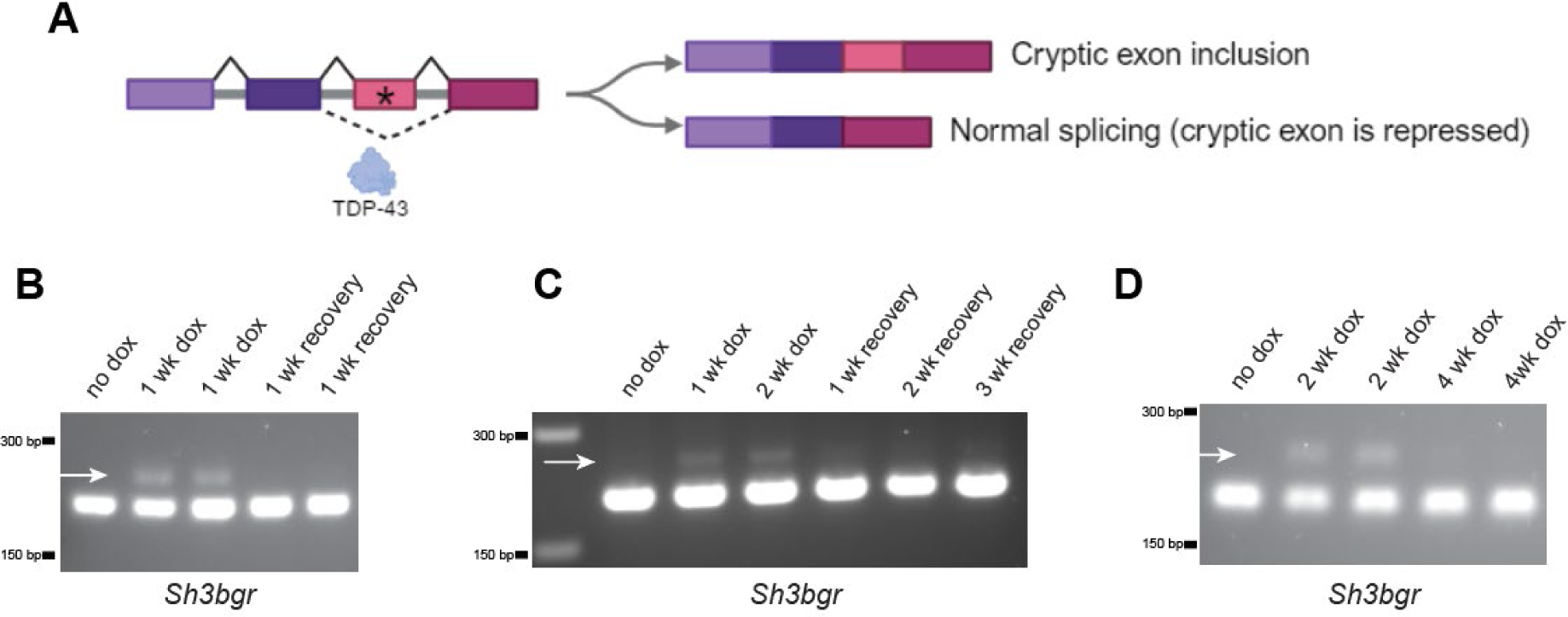
Cryptic exon inclusion is an early event in TDP-43 proteinopathy development. (**A**) Disrupted TDP-43 RNA processing leads to the inclusion of cryptic exons which are normally repressed by TDP-43. (**B**) Agarose gel electrophoresis of products from the RT-PCR amplification of exons 1-2 of *Sh3bgr* from TA muscles of HSA-hTDP-43^ΔNLS^ mice treated for one week with doxycycline and one week after a return to regular chow. A more slowly migrating band is consistent with exon inclusion. (**C**) A similar experiment as in B using mice treated or recovered for the indicated times. (**D**) A similar experiment as in B using mice treated out to four weeks on dox chow.

### TDP-43 seeding persists in HSA-hTDP43^ΔNLS^ mice

We reasoned that similar to patient muscle with TDP-43 aggregates (**Figure 1**), that muscle from our TDP-43 mouse model may have TDP-43 aggregate seeds. To evaluate this, we isolated the insoluble fraction from gastrocnemius muscles of HSA-hTDP43^ΔNLS^ mice and assessed seeding capacity using the TDP-43 biosensor. We found robust seeding from the dox-treated mouse muscle that was not present from no dox controls (**Figure 7A**). We also added the muscle lysate to an α-synuclein FRET sensor line to determine if the lysates were inducing aggregation specific to TDP-43 or if other aggregation-prone proteins could be affected. There was no seeding activity in the α-synuclein biosensors (**Figure 7A**) indicating a TDP-43 specific process. In both assays, seeding was compared to positive controls of recombinant pre-formed fibrils of TDP-43 and α-synuclein. TDP-43 seeding was also confirmed using a different seeding assay that utilizes a cell line which expresses mcherry-tagged TDP-43^ΔNLS^ [31]. Insoluble fractionation western blots of these cells treated with insoluble fractions of 4 week on dox HSA-hTDP43^ΔNLS^ mouse muscle showed increased high molecular weight smears when blotted for TDP-43, consistent with templated aggregate conversion of the soluble mcherry-tagged TDP-43^ΔNLS^ (**Figure 7B**). Most surprisingly, despite no obvious aggregates via immunostaining and western blot at the 3 week recovery timepoint (**Figure 4**), seeding as detected with the TDP-43 biosensor peaked at 1 week of recovery and was still prominent after 3 weeks of recovery (**Figure 7C**). To follow up on how long TDP-43 seeds persist in mouse muscle after the transgene is turned off, a second cohort of mice were treated with dox chow for 2 weeks and collected at recovery timepoints out to 8 weeks at which point seeding had decreased but was still present (**Figure 7D**). These results suggest that TDP-43 aggregate seeds persist in skeletal muscle when obvious signs of TDP-43 aggregation, proteostatic dysfunction or alterations in TDP-43 cryptic splicing are absent.

**Fig. 7.**
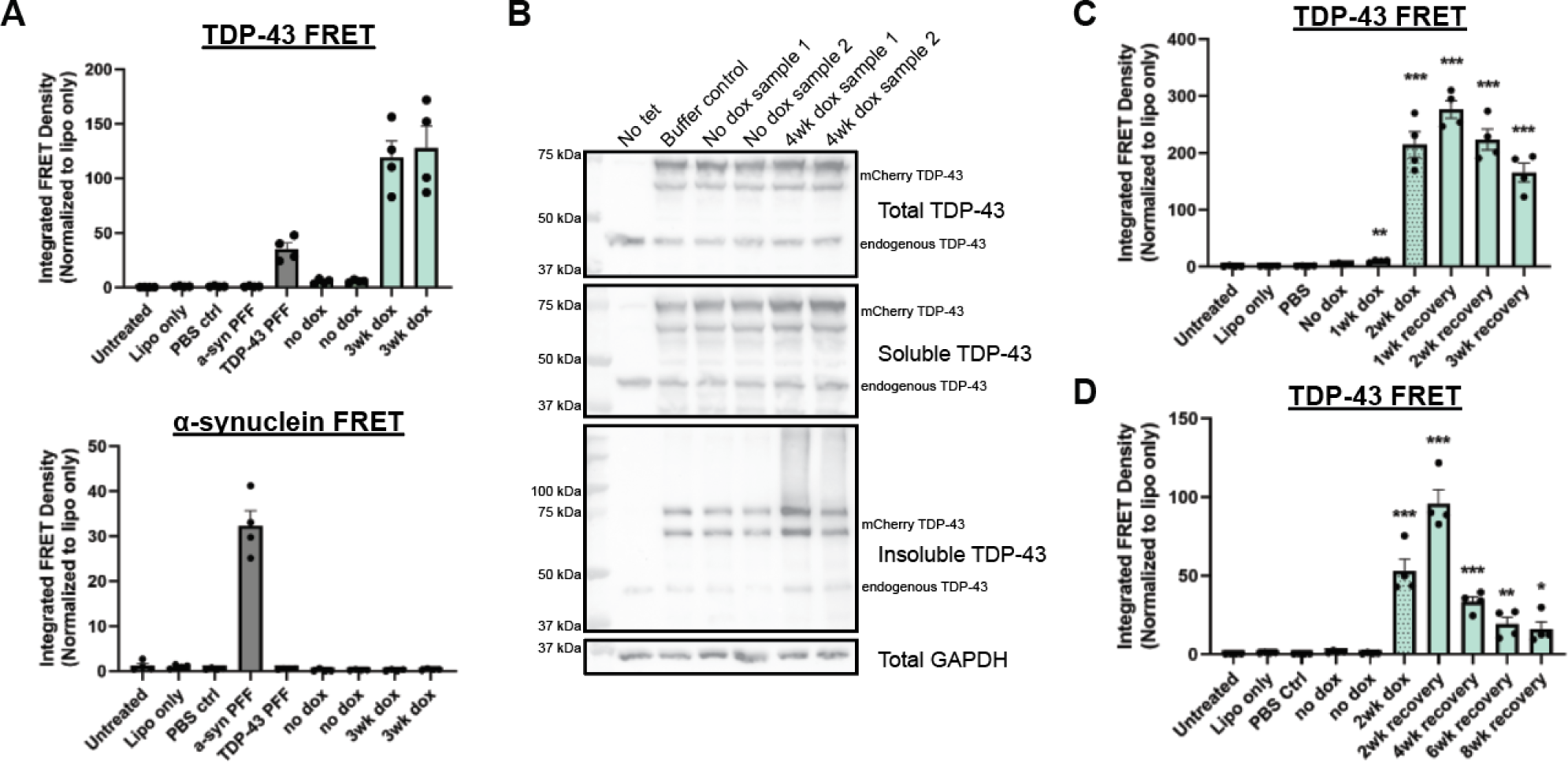
TDP-43 seeding in HSA-hTDP-43^ΔNLS^ mouse muscle is detected by FRET assay. (**A**) The insoluble fractions from muscle of HSA**-**hTDP43^ΔNLS^ mice either treated with no dox or 3 weeks of dox chow were applied to the TDP-43 biosensor and an α-synuclein biosensor. Recombinant TDP-43 and α-synuclein PFF were used for positive controls for each cell line. The insoluble fractions from mice treated with dox chow for three weeks showed robust seeding in the TDP-43 biosensor and none in the α-synuclein biosensor, indicating aggregate specificity. (**B**) Immunoblot from HEK293 cells expressing a tetracycline-inducible mcherry-tagged TDP-43^ΔNLS^. Expression of TDP-43^ΔNLS^ was induced in the cell line for 16 hours and the insoluble fraction from 4 week on dox HSA**-**hTDP43^ΔNLS^ mouse muscle was added to the cells and incubated for 72 hours. The cell lysates were then prepared for insoluble fractionation western blot. The smear of insoluble protein in the lanes of 4 week dox treated muscle indicates successful aggregate seeding. (**C**) HSA-hTDP43^ΔNLS^ mice were treated with dox for two weeks and allowed to recover with regular chow for three weeks, as in **Figure 4**. Seeding peaked 1 week following the return to dox chow and persisted by three weeks. (**D**) Follow-up study demonstrating recovery out to 8 weeks where seeding was still present.

### TDP-43 seeding negatively correlates with histological indications of TDP-43 pathology

Given the discrepancy between visible aggregates and presence of TDP-43 seeding in our mouse model, we next wanted to compare TDP-43 aggregation and seeding in a larger set of human patients. This time, muscle biopsies from 24 patients with sporadic inclusion body myositis were compared to 5 healthy controls, 10 patients with immune-mediated necrotizing myopathy (IMNM), and 10 patients with ALS (6 sporadic, 2 C9ORF72, 1 SOD1, 1 VCP) (**Figure 8A**). Of the 24 IBM samples, 14 had significant TDP-43 seeding. The average integrated FRET density of IBM patient samples was significantly higher than healthy controls (**Figure 8B**). When analyzed using Fisher’s exact test, this gives a sensitivity of 69% and a specificity of 82% with a p value of 0.0001. Next, the amount of FRET seeding activity was compared with histological findings for each patient. Interestingly, there was a negative correlation between the amount of FRET seeding and the percentage of fibers with obvious TDP-43 aggregates or rimmed vacuoles (**Figure 8C-D**). **Figure 8E** shows representative images of a disease control without TDP-43 aggregates, a patient with prominent large TDP-43 aggregates but low FRET seeding, and a patient with a small percentage of fibers with TDP-43 aggregates but high FRET seeding (each indicated by the arrows in **Figure 8A**). This data further supports the presence of TDP-43 prion-like seeds in patient skeletal muscle and that there appears to be a difference between TDP-43 aggregates and seeds that may be difficult to resolve with immunostaining. There did not appear to be any significant relationship between amount of FRET seeding and other clinical data available in **Supplemental Tables 1-2**.

**Fig. 8.**
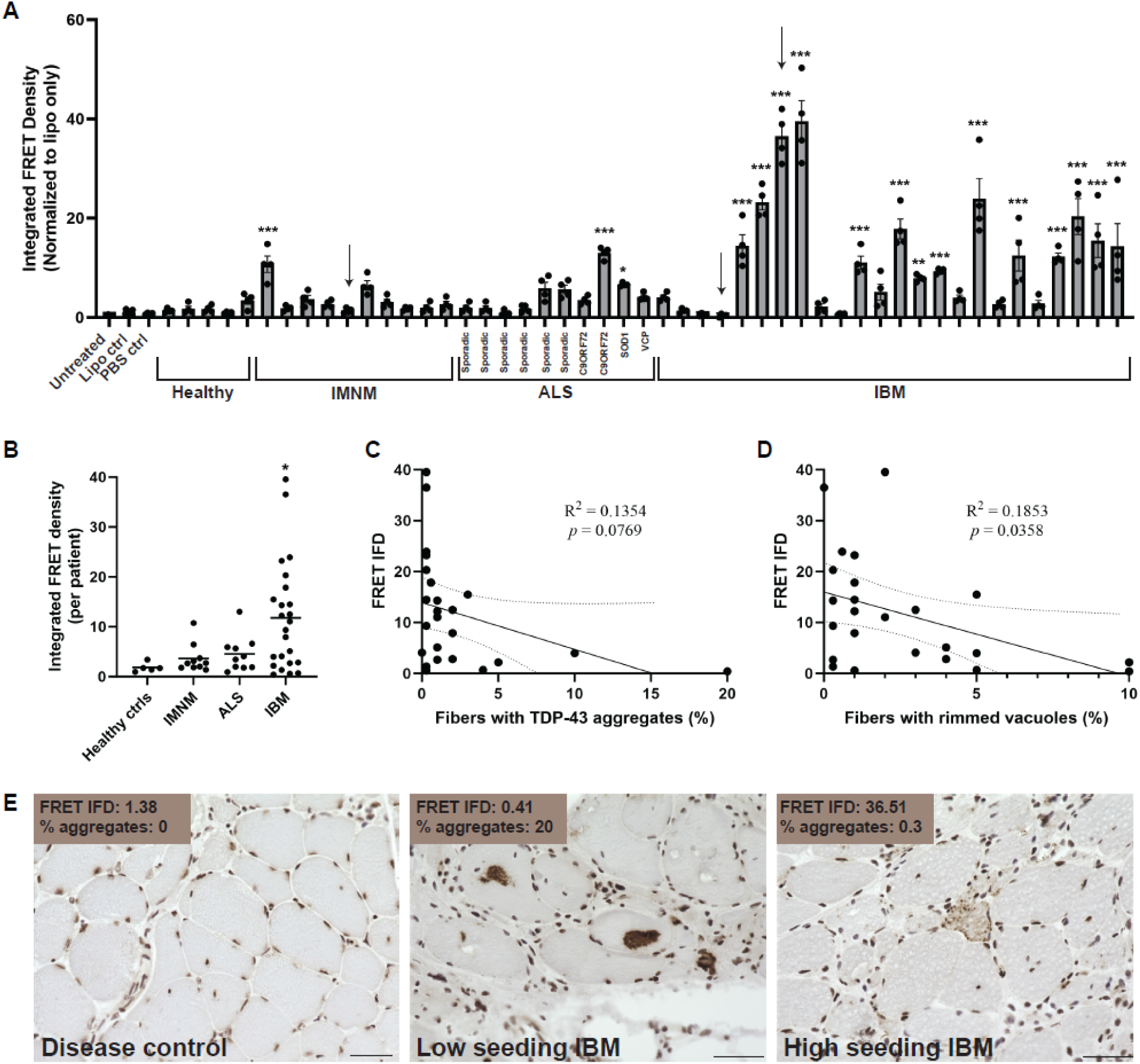
TDP-43 seeding is present in sIBM patient muscle but does not correlate with histological pathology. (**A**) The insoluble fraction from muscle biopsies of 24 sIBM patients, along with 5 healthy controls, 10 with IMNM, and 10 with ALS were added to the TDP-43 FRET sensor line. Of the sIBM samples, 14 were statistically significant. *P<0.05, **P<0.01, ***P<0.001 One-way ANOVA with Dunnett’s multiple comparisons test to lipofectamine only control. (**B**) Representation of each patient’s average integrated FRET density (IFD). P*<0.05 with one-way ANOVA and Dunnett’s multiple comparisons test to healthy controls. The amount of seeding activity had a negative correlation with histological hallmarks of IBM such as TDP-43 aggregates (**C**) and rimmed vacuoles (**D**). (**E**) Representative images of a disease control, low seeding sample, and high seeding sample with their corresponding FRET values (samples indicated by arrows in (**A**). Scalebars represent 50 µm.

## DISCUSSION

Prion-like propagation of aggregation-prone proteins such as tau, α-synuclein and more recently TDP-43 has been identified in the context of neurodegeneration within the central nervous system [1]. Our data supports that TDP-43 in skeletal muscle behaves similarly to mechanisms of neurodegeneration. Specifically, we find that both human patient and experimental models of TDP-43 myopathies have aggregate seeds that are capable of templating the aggregate conversion of monomeric TDP-43 in biosensor cell lines. To date, the presence of aggregate seeds or propagation of aggregate proteins in skeletal muscle has not been demonstrated, with the exception of prion protein in bovine spongiform encephalopathy, scrapie, and chronic wasting disease [19–21].

To study the relationship between TDP-43 aggregation and seeding *in vivo*, we used a tet-on conditional expression system to create a mouse model of TDP-43 aggregation specific to skeletal muscle which can be temporally controlled to study both the accumulation and clearance of TDP-43 aggregate pathology. These mice show significant accumulation of insoluble sarcoplasmic TDP-43 aggregates that result in disrupted proteostasis and alterations in RNA processing as indicated by cryptic exon splicing changes. This data agrees with recent studies from Britson et al. that identified widespread TDP-43 dysfunction in IBM patient muscle biopsies by amplifying transcripts for cryptically included exons [32]. This finding suggests that TDP-43 loss of function may underlie the pathogenesis of IBM. Our data supports a model in which nuclear TDP-43 moves to the sarcoplasm (perhaps during myogenesis or other stresses) and aggregates. Sarcoplasmic aggregates sequester TDP-43, creating a loss of TDP-43 function and cryptic splicing. In our mouse model, TDP-43 splicing dysfunction occurs acutely and resolves.

One intriguing finding from the mouse model is the emergence of TDP-43 aggregate seeds during the clearance phase of TDP-43 inclusions. Specifically, TDP-43 seeding increases following one week of inhibiting transgene expression and persists up to 8 weeks when insoluble TDP-43 protein and myopathologic features of aggregate accumulation have resolved. It is conceivable that large insoluble TDP-43 inclusions are unable to serve as seeds for further TDP-43 aggregates. This would not be unlike studies in yeast in which the yeast prions RNQ1 and Sup 35 must be fragmented into smaller propagons, increasing the seeding capacity [33]. Similarly, a recent study suggested that TDP-43 needed to be proteolytically cleaved prior to its ability to template the aggregation of monomeric TDP-43 *in vitro* [34].

The remodeling of TDP-43 inclusions is likely a dynamic process that requires chaperones and proteases. Notably, the formation and resolution of TDP-43 aggregates or myo-granules may be a normal cellular process in skeletal muscle [22]. Degeneration and regeneration of myocytes not only occurs in pathologic conditions but also occurs during normal muscle growth and repair. Whether TDP-43 seeds are present and propagate under these conditions is not known. Therefore, additional models of inducing skeletal muscle injury and regeneration to trigger physiologic TDP-43 myo-granule formation and potentially seeding from endogenous TDP-43 are also being considered. These include *in vitro* cell stress assays by arsenite treatment, *in vivo* barium chloride injections to induce muscle degeneration and regeneration, or eccentric contraction muscle injury models. By introducing myopathy-causing gene mutations into these systems we can examine how these mutated proteins might be influencing TDP-43 seeding behavior. Inherited myopathies with TDP-43 inclusions are associated with mutations in TDP-43 itself but mutations in other myopathy-linked genes have been shown to influence TDP-43 granule formation and resolution. For example, dominant mutations in *VCP* cause multisystem proteinopathy, a degenerative syndrome characterized by inclusion body myopathy, ALS and FTD. The three diseases are unified pathologically by TDP-43 pathology. VCP is necessary for the clearance of cytosolic stress granules and may lead to the persistence of TDP-43 granules in MSP patients [25, 35]. Other muscle-derived TDP-43 proteinopathies include Welander distal myopathy that is associated with the stress granule protein TIA1 [36]. TIA1 mutations lead to an increase in stress granule formation and decreased clearance [37]. These defects may contribute to the development of TDP-43 pathology.

While TDP-43 aggregates are a marker of sporadic inclusion body myositis with between 78 to nearly 100% of IBM biopsies having at least one aggregate containing fiber, the burden of these aggregates can be quite variable ranging from 0.5% to 25% of fibers containing aggregates [8, 9]. This raises questions regarding the relevance of TDP-43 aggregation or dysfunction as the primary driver of disease pathogenesis in IBM. In this study, we used a TDP-43 aggregate biosensor to detect TDP-43 seeding from IBM patient muscle biopsies. Interestingly, the presence of seeding had a negative correlation with the amount of TDP-43 aggregate staining per patient. These data are consistent with our findings in the mouse model in which TDP-43 seeding is still present despite the clearance of larger aggregates. Our studies support that a pathogenic TDP-43 aggregate species that is not seen by traditional immunohistochemical stains is present. As we have demonstrated seeding capability of muscle-derived TDP-43 from both mouse and humans, we hypothesize that skeletal muscle could potentially serve as a reservoir for TDP-43 seeds as it does for the prion protein.

## MATERIALS AND METHODS

### Biosensor culture and seeding

The TDP-43 fluorescence resonance energy transfer (FRET) sensor lines were received from Marc Diamond lab [26]. These cells express aa262-414 of TDP-43 with c-terminal mruby3 or mclover3 fluorescent tags. Cells were maintained in 10% FBS medium and split using 0.05% trypsin. For detecting TDP-43 seeding in cell or tissue lysates, the FRET cells were plated onto a 96-well plate at a density of 35,000 cells/well. After about 24 hours, the cells were around 50% confluent and treated with seeds. Recombinant seeds or insoluble fractions resuspended in PBS were added to the cells along with Lipofectamine 2000 transfection reagent (1uL lipofectamine per well; Thermo Fisher, 11668019). Cells were left to incubate and fixed after 72 hours. To fix, cells were trypsinized and centrifuged at 1000 xg for 5 minutes in a round-bottom 96w plate, then resuspended in 2% PFA for 10 minutes, then centrifuged again and resuspended in 150uL of flow running buffer and stored at 4°C until ready for use.

The cells were analyzed by flow cytometry using the MacsQuant VYB. The mclover3 was excited by a 488 nm laser and detected with a 525/50 nm bandpass filter. The mruby3 signal was excited by a 561 nm laser and detected with a 615/20 nm filter. FRET signal was excited by a 488 laser and detected with a 614/50 nm filter. The integrated FRET density was calculated as % positive cells multiplied by the median fluorescence intensity and normalized to a buffer only control.

As a further confirmation of TDP-43 seeding *in vitro*, insoluble fractions were added to HEK293 cells stably expressing tetracycline-inducible mcherry-tagged TDP-43^ΔNLS^ [31]. These cells express soluble TDP-43^ΔNLS^ when activated with tetracycline but can be induced to aggregate with the addition of stressors or TDP-43 seeds. Expression was induced with 1 µg/mL tetracycline 16 hours before the addition of insoluble fraction. After 72 hours of incubation with the insoluble fraction, aggregation was measured by fractionation western blot for insoluble TDP-43.

### Formation of recombinant TDP-43 and α-synuclein pre-formed fibrils (PFF)

Recombinant TDP-43 monomer and pre-formed fibrils were generated as previously described [38, 39]. BL21(DE3)-RIL *Escherichia coli* cells were transformed with pJ4M-TDP43-MBP-His (Addgene 104480) and grown on LB/Kanamycin/Chloramphenicol plates then used to inoculate 2L of TB/Kan/Cam/2g dextrose. Expression of TDP-43 was induced at OD_600_ = 0.5 with addition of 1 mM IPTG for 18 hours at 16°C. Cell pellets were resuspended in 50 mM HEPES, pH 7.4, 0.5 M NaCl, 30 mM imidazole, 10% glycerol, 2 mM DTT, 100 µM PMSF, 10 µM Pepstatin A, supplemented with lysozyme. Lysates were sonicated at 40% amplitude, 30s on/30s off for 3 cycles. Lysates were purified through binding to Fast flow Nickel sepharose (GE Healthcare) and washed with buffer (50 mM HEPES, 10% glycerol, 0.5 M NaCl, 30 mM Imidazole, pH=8, 2 mM DTT) and eluted in elution buffer (50 mM HEPES, 10% glycerol, 0.5 M NaCl, 0.5 M Imidazole, pH=8, 2 mM DTT, 0.1 mM PMSF, 10 µM PepstatinA, cOmplete EDTA-free protease inhibitors). The resulting eluent was further purified with amylose resin (New England BioLabs E8021S) which binds MBP and then washed with amylose wash buffer (50 mM HEPES, 10% glycerol, 0.5 M NaCl, 2 mM DTT) and eluted with amylose elution buffer (50 mM HEPES, 10% glycerol, 0.5 M NaCl, 2 mM DTT, 0.1 mM PMSF, 10 µM pepstatin A, 10 mM Maltose, and cOmplete EDTA-free protease inhibitors). Protein concentration was determined by Bradford assay and samples were concentrated to approximately 40 µM then flash frozen and stored at −80 °C until use. To induce protein aggregation, 10 µM of TDP-43 monomer was diluted in assembly buffer (50 mM HEPES, pH 7.4, 10% glycerol, 1 mM DTT), and shake in an Eppendorf Thermomixer for 1 week at 30°C and 650 rpm. Aggregation reactions were initiated by addition of TEV protease to cleave off the MBP-His tag and turbidity was monitored by continuously measuring absorbance at 395 nm at 30°C with agitation over 12 hours in a BioTek Epoch plate reader.

Recombinant α-synuclein preformed fibrils were used as a positive control with the α-synuclein FRET sensor line when testing samples for specificity to TDP-43 aggregation only. The recombinant α-synuclein was created as previously reported [40, 41]. Briefly, the recombinant protein was extracted from *E. coli* cells by osmotic shock and purified using heat precipitation and ion exchange chromatography with diethylaminoethyl resin. The purified protein was dialyzed overnight into 10 mM Tris-HCl, pH 7.6, 50 mM NaCl, 1 mM DTT was dialized overnight and protein yield was determined by BCA assay and SDS-PAGE. To form fibrils, 2 mg/mL of α-synuclein monomer was diluted in 20 mM Tris-HCl, pH 8.0, 100 mM NaCl and shaken in an Eppendorf Thermomixer for 72 hours at 37 °C and 1000 rpm. To obtain the concentration of fibrils, the protein was centrifuged at 18,000xg for 15 minutes to separate monomer from fibril and the concentration of monomer in the supernatant was measured by BCA assay and used to calculate the fibril concentration.

### Insoluble fractionation

Cell lysates were collected in radioimmunoprecipitation assay (RIPA) buffer with protease inhibitor cocktail (Sigma-Aldrich, S8820). Muscle and brain samples were also lysed with RIPA buffer during a homogenization step using mortar and pestle. The samples were left to sit for at least one hour with frequent vortexing, then centrifuged at 13,000 rpm for 10 minutes at 4 °C to pellet the tissue debris. The concentration of the supernatant was determined by BCA assay and each sample was diluted to equal amounts of starting protein before beginning the fractionation process. Samples were sonicated with 10 cycles of 30s on, 30s off at amplitude 50 and then the insoluble fraction was pelleted through ultracentrifugation at 100,000 xg for 30 minutes. If preparing samples for western blot, the pellet was resuspended in a RIPA wash, sonicated again and ultracentrifuged again. The final pellet was resuspended in 7M urea buffer with 2M thiourea, 4% CHAPS, and 30mM Tris pH 8.5. If the sample was prepared for inducing seeding in a cell line, the initial pellet was washed with PBS for the second sonication and ultracentrifugation step. The final pellet is then resuspended in PBS.

### Generation of HSA-TDP43^ΔNLS^ mice

To create a muscle-specific doxycycline-inducible mouse model of TDP-43, HSA-rtTA mice (The Jackson Laboratory, strain 012433) were crossed with tetO-hTDP-43^ΔNLS^ mice (The Jackson Laboratory, strain 014650). This created a tet-on expression system in which TDP-43^ΔNLS^ was expressed under the *ACTA1* skeletal muscle promoter only when mice received a doxycycline (dox) chow diet (200 mg/kg, Fisher Scientific, 14-727-450). As controls, WT C57BL/6, single transgenic (tetO-hTDP-43^ΔNLS^), and double transgenic mice with no dox treatment were compared to double transgenic with dox treatment. All mice used in this study were bred on a C57BL/6 background and genotyped by Transnetyx using the primers found in **Table 1**. Male and female mice were used, without any noted differences. Animal housing and procedures were in accordance with protocols approved by the Animal Studies Committee at Washington University in St Louis.

**Table 1:**
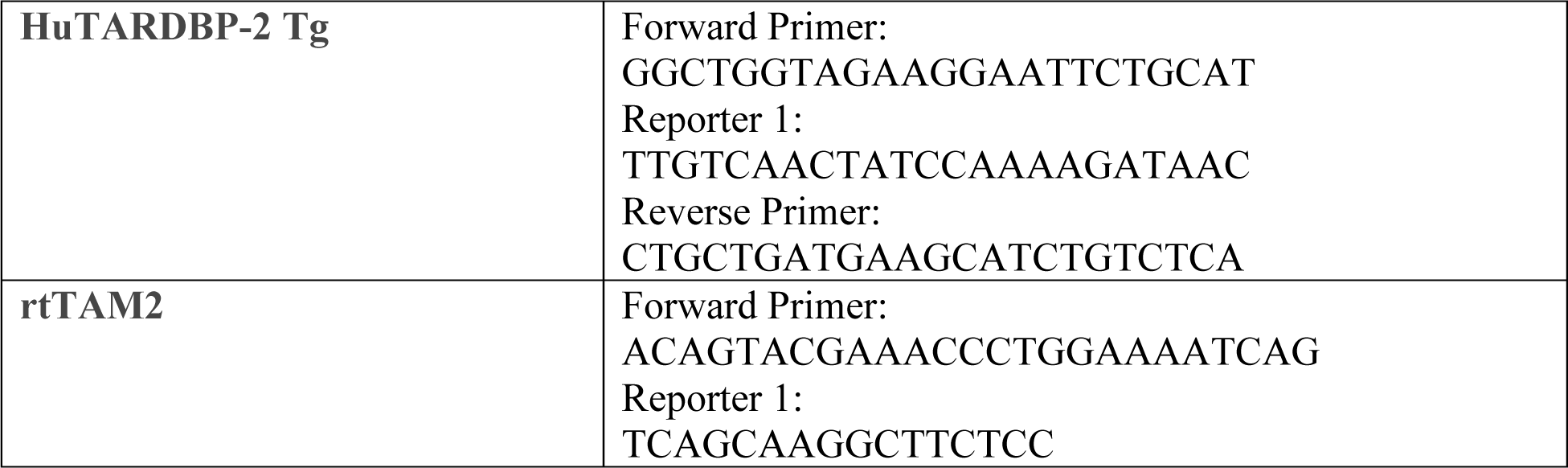
mouse genotyping primers.

### Mouse strength analysis

Grip strength was measured using a trapeze bar attached to a force transducer to record peak-generated force (Stoelting Co.). The mice were placed with forepaws gripping the bar and were pulled backwards by the tail until unable to maintain their grip. This maximum force generation was recorded. Each mouse was tested 5 times and the highest and lowest values were dropped and the remaining three were averaged and divided by mouse weight. For the hanging wire test, mice were placed in container with a mesh grid bottom. Once the mouse was settled on all fours and gripping the mesh bottom, the container was inverted, 15cm above a cage. The amount of time before the mouse released their grip on the mesh was recorded, and the average of three trials was again divided by the mouse weight. For both tests, mice were cycled through to allow each mouse to rest between trials.

### X-ray imaging and kyphosis analysis

After three weeks of dox chow, mice were anesthetized and imaged using a Faxitron UltraFocus100 x-ray machine with VisionDXA imaging software. The kyphosis index was calculated according to the protocol in Laws and Hoey 2004 [42]. Briefly, line AB is drawn from vertebrae C7 to L6 and line CD was drawn from line AB to the farthest vertebral body. Kyphosis index was calculated as the length of line AB divided by the length of line CD.

### Human brain and muscle tissue

Postmortem human motor cortex and psoas muscle from was collected from ALS patients who consented for autopsy with a post-mortem interval that ranged from 7-48 hours. Tissues were flash frozen in liquid nitrogen prior to storage at −80C. The presence of TDP-43 pathology was confirmed by clinical neuropathological analysis in sALS and *C9ORF72* fALS cases. The use of muscle biopsies in **Figure 1** was approved by the Washington University in St Louis Institutional Review Board (IRB201104149). The use of human sIBM muscle biopsy samples in **Figure 8** was approved by Johns Hopkins Institutional Review Board (IRB0072691). De-identified clinical information is available in **Supplemental Tables 1-2**.

### Histochemistry/Immunohistochemistry

Skeletal muscle samples were mounted in tragacanth gum (10% solution, Sigma-Aldrich, G1128) and flash frozen in 2-methylbutane over liquid nitrogen and stored at −80 °C until ready to be sectioned to 10 micron thickness. For hematoxylin and eosin (H&E) stain, a 1% aqueous solution of eosin Y (Sigma E-6003) was prepared in deionized water and Harris Hematoxylin stain (Lerner Laboratories 1931382) was filtered before use. Slides in a metal staining rack were immersed in the filtered Harris Hematoxylin for 10 seconds then transferred to a beaker of tap water and rinsed until the water is clear. Then the slides were immersed in eosin stain for 30 seconds and again rinsed with tap water. Then sections were dehydrated in ascending alcohol solutions (50%, 70%, 80%, 95%x2, 100%x2), cleared with xylene 3-4 times, and a glass coverslip was mounted to the glass slide using Permount.

For nicotinamide adenine dinucleotide diaphorase (NADH) staining, NADH solution (80mg of NADH in 50mL of 0.05M TRIS buffer, pH7.6) was combined 1:1 with nitro-blue tetrazolium (NBT) solution (200mg in 100mL of TRIS buffer). Slides were incubated in the 1:1 solution for 30 minutes at 37 °C then washed 3 times with water. Unbound NBT was removed by three exchanges each in 30%, 60%, and 90% acetone solutions, increasing then decreasing concentration. Finally, slides were rinsed several more times in water then mounted with a coverslide using Permount.

For TDP-43 staining of human muscle samples, the sections were fixed for 20 minutes in 4% paraformaldehyde, washed with PBS for 10 minutes then blocked and permeabilized with 0.1% Triton-X in PBS and 5% normal goat serum for 1 hour at room temperature. The primary antibody was diluted in blocking buffer and incubated overnight at room temperature. On the second day, slides were rinsed 3x 10 minutes with 0.1% PBS-T then incubated with 30% H2O2 for 15 minutes at room temperature. The biotin-labeled secondary antibody was diluted in blocking buffer and incubated for 1 hour in a humid chamber then washed 3x 10 minutes with PBS-T. The slides were then incubated with ABC reagent (Vectastain kit) for 60 minutes in the humid chamber and again washed 3x 10 minutes with PBS-T. DAB (Vectorlabs SK-4100) was added to the slides to develop and the reaction was quenched with water and rinsed in water for 5-10 minutes. Sections were dehydrated with 4 changes of 95% ethanol for 30 seconds each, 4 changes of 100% ethanol for 30 seconds each, and 4 changes of xylene for 2 minutes each before mounting the coverslip.

Sections were fixed for immunostaining using 3.7% paraformaldehyde for 10 minutes followed by 10 minutes of ice-cold acetone. The muscle sections were then permeabilized for 20 minutes in 0.5% triton and blocked for 2 hours at RT in Perkin Elmer blocking reagent (FP1012). Primary antibodies were diluted in blocking reagent according to dilutions listed in **Table 2** and incubated at 4°C overnight or for 1 hour at room temperature. After 3 rinses for 5 minutes each with 1x PBS, secondary antibodies were added to the slides at 1:1000 dilution in blocking reagent and incubated for 1 hour at room temperature. Slides were rinsed with 1x PBS again 3 times for 5 minutes each, then incubated for 10 minutes with 1 µg/mL DAPI followed by a final 3 PBS rinses. Cover glass was mounted to slides using Mowiol 4-88 (Sigma Aldrich, 81381).

**Table 2:** antibody dilutions used for immunocytochemistry/immunohistochemistry.

| **Marker** | **Catalog #** | **Dilution** |
| --- | --- | --- |
| TDP-43 | Proteintech 10782-2-AP | 1:1000 (mouse tissue)<br>1:2000 (human tissue) |
| pTDP-43 | Biologend 829901 | 1:1000 |
| P62 | Abcam AB118275 | 1:1000 |
| Poly-ubiquitin | Dako Z0458 | 1:200 |

### Imaging and image analysis

Stained mouse muscle sections were imaged using a Nikon Eclipse 80i microscope and MetaMorph Imaging Series 7.8. Human muscle staining was imaged with a Keyence BZ-X710. To calculate the percentage of human muscle fibers positive for TDP-43 aggregates, 2-3 images of different 20x fields of view were taken per biopsy and around 300 fibers were counted.

### Western blotting

Cell or muscle tissue lysates were collected in RIPA buffer with protease inhibitor cocktail. Protein concentrations were obtained through BCA assay and measured with the BioTek Epoch plate reader at 562 nm. Each lane of a 10% SDS-PAGE gel was loaded with 20-40 µg of total protein or equal volumes of soluble or insoluble samples, and gels were run at 100 V for 10 minutes followed by 120 V for 1 hour 15 minutes. Next, the samples were transferred to nitrocellulose membranes at 26 mA for 2 hours. Membranes were blocked with 5% milk for 1 hour then primary antibodies were added according to the dilutions listed in **Table 3** and incubated overnight at 4°C. Blots were treated with anti-rabbit or anti-mouse HRP secondary antibodies at a 1:5000 dilution in 5% milk for 1 hour at room temperature then rinsed 3 times with 1x PBS-tween before imaging membranes. Membranes were imaged using ECL (Bio-rad, 1705060) and Syngene G:Box Chemi XT4 imager.

**Table 3:** antibody dilutions used for western blotting.

| **Marker** | **Catalog #** | **Dilution** |
| --- | --- | --- |
| TDP-43 | Proteintech 10782-2-AP | 1:1000 |
| pTDP-43 | Proteintech 22309-1-AP | 1:1000 |
| GAPDH | Cell Signaling 2118 | 1:1000 |
| Poly-ubiquitin | Enzo ABS840 | 1:1000 |
| P62 | Abnova H0008878 | 1:1000 |
| VCP | BD 612182 | 1:500 |
| optineurin | Proteintech 10837-1-AP | 1:2000 |
| LC3 | Sigma L7543 | 1:500 |
| Lamp2 | Invitrogen PA1-655 | 1:500 |
| HSP70 | Enzo ADI-SPA-810 | 1:500 |
| Dnajb6 | Abcam ab198995 | 1:500 |

### Cryptic exon detection

RNA was extracted from tibialis anterior muscles by homogenization in Tri Reagent (Molecular Research Center, Inc) according to the reagent protocol. Total RNA was converted to cDNA using the High Capacity cDNA Reverse Transcription Kit (Applied Biosciences, 4374966). Primers for *Sh3bgr* cryptic exon detection (**Table 4**) were used to amplify the targets using the DreamTaq 2x master mix (Thermo Fisher, K1081). The PCR reaction settings using a Bio-Rad C1000 Touch Thermal Cycler were 95 °C for 60s, 40 cycles of 95 °C for 30s, 63 °C for 15s, 72 °C for 45s, then 7 minutes at 72 °C before holding at 4 °C. Amplified samples were run on a 2% agarose gel (Lamda Biotech, Inc., A113-3) containing 0.5μg/mL ethidium bromide (Millipore Sigma, E1510) for 1.5 hours.

**Table 4:**
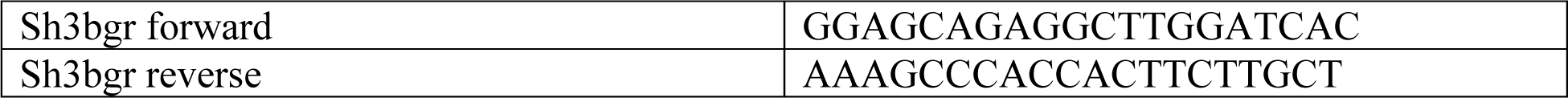
primers used for RT-PCR to detect cryptic exons.

### Transmission electron microscopy

Mice were anesthetized and perfused with warmed Lactated Ringer’s Irrigation solution (B. Braun Medical Inc) followed by fixative solution (2.5% glutaraldehyde, 2% paraformaldehyde, 0.15M cacodylate buffer pH 7.4 with 2 mM CaCl_2_). Tibialis anterior muscles were dissected and fixed overnight at 4 °C in the same fixation solution. Post fixation, samples were rinsed in 0.15 M cacodylate buffer containing 2 mM calcium chloride 3 times for 10 minutes each followed by a secondary fixation in 1% osmium tetroxide and 1.5% potassium ferrocyanide in 0.15 M cacodylate buffer containing 2 mM calcium chloride for 1 hour in the dark. The samples were then rinsed 3 times for 10 minutes each in ultrapure water and *en bloc* stained with 2% aqueous uranyl acetate overnight at 4 °C in the dark. After 4 washes for 10 minutes each in ultrapure water, the samples were dehydrated in a graded acetone series (10%, 30%, 50%, 70%, 90%, 100% x3) for 10 minutes each step, infiltrated with Spurr’s resin (Electron Microscopy Sciences) and embedded and polymerized at 60 °C for 48 hours. Post curing, 70 nm thin sections were cut and imaged on a TEM (Jeol JEM-1400 Plus) at 120 kV.

### Statistical Analysis

All quantitative data was analyzed in GraphPad Prism. Graphs are presented as mean ±SEM. Experiments involving the FRET sensor line included 4 replicate wells treated with the same insoluble sample or other control condition. One-way ANOVA followed by Dunnett’s multiple comparisons test was used to detect significance compared to the lipofectamine only control. When comparing the average of only two groups such as the no dox versus dox-treated mouse muscle weights, a two-tailed unpaired Student’s *t* test was used. In all cases: *P>0.05, **P<0.01, ***P<0.001.

## Acknowledgements

We thank the Marc Diamond lab of University of Texas Southwestern for the generating and sharing the TDP-43 FRET biosensor line used in these studies.

We acknowledge the assistance of John Wulf II and Gregory Strout at the Washington University Center for Cellular Imaging (WUCCI) in electron microscopy studies, which is supported by Washington University School of Medicine, The Children’s Discovery Institute of Washington University and St. Louis Children’s Hospital (CDI-CORE-2015-505 and CDI-CORE-2019-813) and the Foundation for Barnes-Jewish Hospital (3770 and 4642).

## Author contributions

E.M.L. designed and carried out experiments and analyzed results. S.P. coordinated animal husbandry for generating the mouse model. J.D. optimized PCR protocol for detecting cryptic exons and sectioned muscle samples for histology. D.D.D., P.K., S.C., and M.E.J. provided monomeric and PFF forms of recombinant α-synuclein and TDP-43 for controls in FRET experiments. Y.M.A. contributed the TDP-43-mcherry cell line for an additional aggregation assay. C.V.L. provided ALS patient tissue, and C.I., T.E.L. and A.P. provided IBM patient muscle tissue. C.I. conducted staining and analysis of IBM patient muscle tissue. E.M.L. and C.C.W. wrote the manuscript which was edited and approved by all authors.

## Data and materials availability

All data are available in the manuscript or supplemental files. The mouse model was generated from two mouse lines available for purchase from The Jackson Laboratory.

